# Gene editing in Farm Animals: A Step Change for Eliminating Epidemics on our Doorstep?

**DOI:** 10.1101/2021.04.19.440533

**Authors:** Gertje Eta Leony Petersen, Jaap Buntjer, Fiona S. Hely, Timothy John Byrne, Bruce Whitelaw, Andrea Doeschl-Wilson

**Affiliations:** AbacusBio Ltd, Dunedin 9016, New Zealand; The Roslin Institute, University of Edinburgh, Roslin Institute Building, Easter Bush EH25 9RG, Scotland, UK; AbacusBio International, Roslin Innovation Centre, The University of Edinburgh, Easter Bush EH25 9RG, Scotland, UK; The Global Academy of Agriculture and Food Security, The University of Edinburgh, Easter Bush EH25 9RG, Scotland, UK

**Keywords:** Gene editing, PRRS, CRISPR/Cas9, mathematical model, infectious disease

## Abstract

Recent breakthroughs in gene-editing technologies that can render individuals fully resistant to infections may offer unprecedented opportunities for controlling future epidemics. Yet, their potential for reducing disease spread are poorly understood as the necessary theoretical framework for estimating epidemiological effects arising from gene editing applications is currently lacking. Here, we develop semi-stochastic modelling approaches to investigate how the adoption of gene editing may affect infectious disease prevalence in farmed animal populations and the prospects and time-scale for disease elimination. We apply our models to the Porcine Reproductive and Respiratory Syndrome PRRS, one of the most persistent global livestock diseases to date. Whereas extensive control efforts have shown limited success, recent production of gene-edited pigs that are fully resistant to the PRRS virus have raised expectations for eliminating this deadly disease.

Our models predict that disease elimination on a national scale would be difficult to achieve if gene editing was used as the only disease control. However, when complemented with vaccination, the introduction of 10% of genetically resistant animals in a fraction of herds could be sufficient for eliminating the disease within 3-6 years. Besides strategic distribution of genetically resistant animals, several other key determinants underpinning the epidemiological impact of gene-editing were identified.

## Introduction

Novel genomic technologies such as gene editing offer promising opportunities to tackle some of the most pressing global challenges humanity faces today. They provide new prospects to solving emerging threats such as the global Covid-19 pandemics (1), as well as to long-standing global health issues such as the HIV/Aids crisis (2) or malnutrition (3, 4), with minimal side effects. Besides the medical field, food production stands to gain most from widespread use of genome editing technologies. Currently 11% of the human population suffers malnourishment (5), and this is expected to increase with the projected growth of the human population to 10.9 billion by 2100 (42%) (6). Meeting the 60% increase of agricultural production needed to provide sustainable and nutritious diets will likely require transformative innovations to existing production methods (7). While genome-editing technologies have been applied widely in plant breeding to simultaneously improve production and resilience to diverse stressors (see (8) for examples), their application in the livestock sector is still in its infancy, primarily due to technical limitations as well as ethical and societal concerns (9). Nevertheless, breakthroughs in genetic modification of farm animals through genome editing start to emerge with drastic improvements in efficiency traits (10, 11), animal welfare (12) and disease resistance (13, 14). Improving disease resistance in livestock seems particularly pertinent, as infectious diseases affect the entire food production chain and its economic viability (15).

The recent scientific breakthroughs in genome editing raise expectations for radical shifts in infectious disease control in livestock (14). They equally evoke a number of pressing questions concerning the theoretical and practical feasibility of tackling diseases for which conventional control methods have failed. It is currently not known how to best implement this novel technology in order to achieve noticeable reduction in disease prevalence and possibly even eliminate the disease on a national scale, and in what time scale such ambitious goals could be achieved.

These questions are impossible to address in an entirely hypothetical context, since epidemiological characteristics affecting the spread of the disease in question and the dynamics of the dispersal of resistant animals within the population play important roles in the success of the scheme. Here, we focus on a particular disease, PRRS, for the development of a mathematical modelling framework to investigate the feasibility of gene editing for disease elimination. PRRS (Porcine Reproductive and Respiratory Syndrome) represents one of the most important infectious disease problems for the pig industry worldwide with economic losses estimated at $2.5 billion per annum in the US and Europe alone (16, 17). Despite extensive global control efforts, the disease continues to persist in national commercial pig populations, largely due to high genetic heterogeneity of the PRRS virus, *PRRSV*, (18) and the associated limited efficacy of all PRRS vaccines and limited reliability of diagnostic tests (19– 21). There is considerable natural genetic variation in pigs’ responses to *PRRSV* infection but evidence to date suggests that no pig is naturally fully resistant to it (22). However, recent advances in gene editing of porcine macrophages, in which a simple disruption of the CD163 gene confers complete resistance to infection with *PRRSV*, may revolutionize future PRRS control (23–25).

To exploit the full potential of gene editing for PRRS control, we here develop a theoretical proof-of-concept model to address a number of crucial questions: To what extent can gene editing help reduce PRRS prevalence in national commercial pig populations? Is it possible to eliminate this disease through gene editing by creating a disease-resistant subpopulation adequately dispersed within the national susceptible population? If so, what proportion of pigs would have to be PRRS resistant and how would these animals need to be distributed across herds?

It is unlikely that gene editing will fully replace existing control methods, such as wide-spread vaccination. Hence, we also use our model to investigate the epidemiological effects of gene editing and vaccination combined. Finally, we investigate how fast the required proportion of resistant animals could be introduced in a national commercial pig population, if gene editing was strictly limited to breeding programs and resistance alleles propagate to commercial pigs using current industry practices. This last question becomes particularly important for an RNA virus with a high evolutionary rate such as *PRRSV*, since escape mutants of the virus might limit the shelf-life of gene editing in terms of effectiveness (14).

We address these questions with two linked simulation models: (1) an epidemiological model to simulate the effects of different disease control schemes on PRRS prevalence in a national commercial pig population, and (2) a gene flow model to simulate the propagation of *PRRSV* resistance alleles from breeding programs that routinely carry out gene editing for PRRS resistance, into the commercial population. The epidemiological model provides insight into the numbers and distribution of genetically resistant pigs required to eliminate PRRS under a range of realistic scenarios. The allele propagation models then provides estimates for the time required to realistically produce this required number of genetically resistant pigs.

Our proof-of-concept model provides the first quantitative estimates for how gene editing may reduce infectious disease prevalence in farm animals and the required time frame and criteria for eliminating a disease on a national level.

## Results

Impact of gene editing on disease prevalence and chance of disease elimination

### Gene editing as the only disease control

We first investigated how gene editing of pigs may affect PRRS prevalence at a national level. We assessed whether disease elimination through gene editing alone is possible and what proportion of a population would have to be genetically resistant to achieve this goal. Epidemiological theory for herd immunity stipulates that disease elimination is possible provided that individual subpopulations or herds contain sufficient proportions of resistant individuals (26). The required proportion of resistant individuals (*P*_*e*_*\**) in a population depends largely on the disease transmission potential, otherwise known as the reproductive ratio *R*, which is defined as the expected number of secondary cases caused by a primary case over its infectious period (27).

To predict the potential effects of gene editing on PRRS prevalence at a national level, we simulated national commercial pig populations consisting of herds that varied in size, PRRS virus exposure and the baseline transmission potential *R*_*0*_ in the absence of genetically resistant or vaccinated animals (see Methods). We then simulated four different distribution scenarios according to which given numbers of available genetically resistant pigs are distributed across the herds. These scenarios mimic different degrees of regulations concerning the distribution of these pigs, ranging from a centrally regulated scheme that may be based on either little or accurate information about the baseline transmission potential *R*_*0*_ to an entirely voluntary uptake by the farmers (see Methods and Table 1). Following epidemiological theory (27), the presence of genetically resistant pigs in a herd reduces the herd specific *R*_*0*-_value to the effective reproductive rate

**Table 1.**
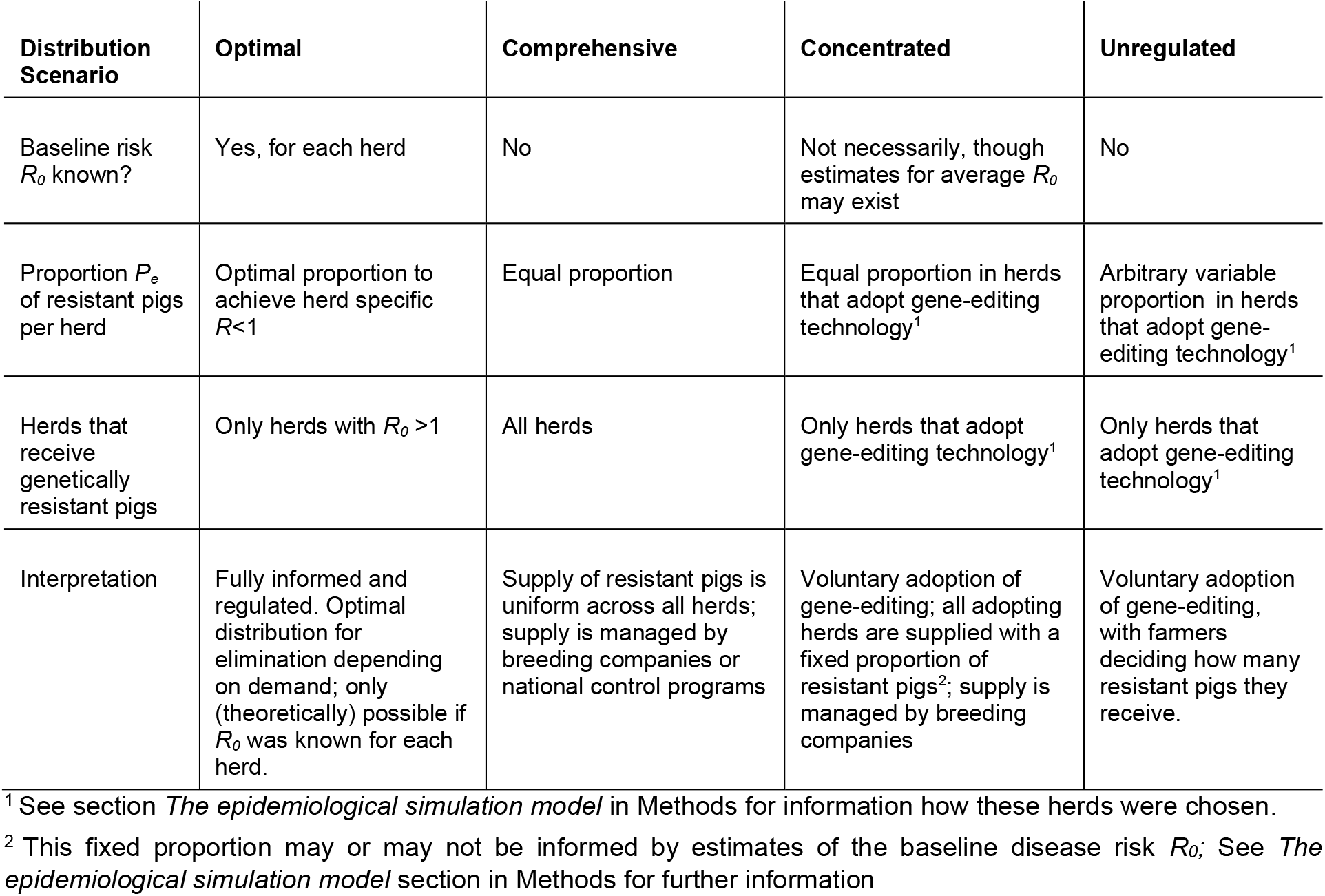
Overview of the scenarios for the distribution of genetically resistant pigs across herds in the epidemiological model.

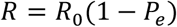

where *P*_*e*_ denotes the fraction of genetically resistant in the herd (see Methods for the more generic model also including vaccination effects). PRRS prevalence on a national level was then defined as the proportion of herds with *R* above one.

Figure 1 demonstrates that gene editing can contribute to considerable reduction of disease prevalence and even lead to full elimination under optimal conditions. However, the rate at which disease prevalence reduces with increasing proportions of genetically resistant individuals depends strongly on both *R*_*0*_ and how resistant pigs are distributed across herds. In particular, the latter plays a significant role in whether or not a strategy achieves full disease elimination (Figure 1).

**Figure 1.**
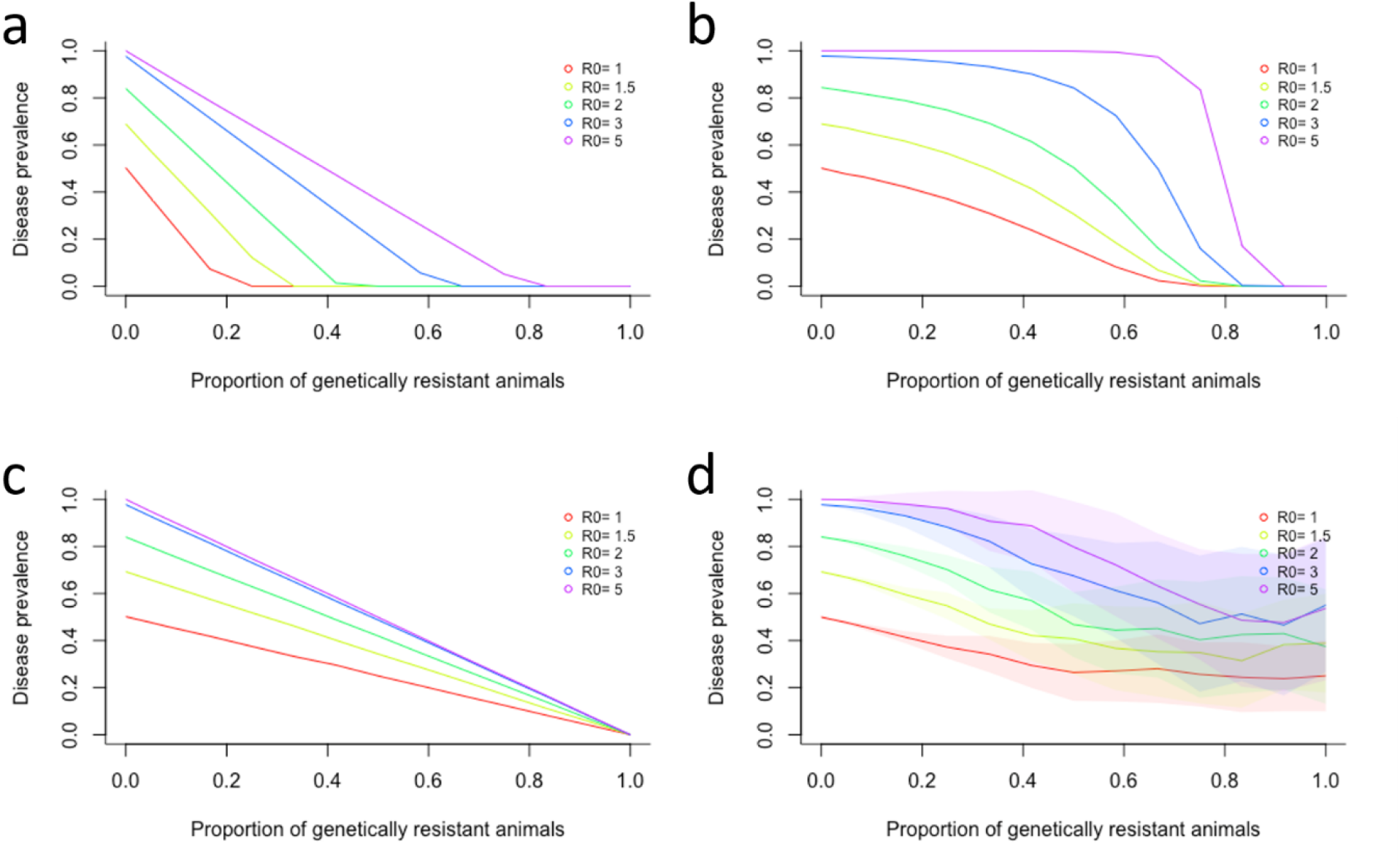
Predicted reduction in PRRS prevalence achieved by using genetically PRRS resistant pigs, depending on the average baseline PRRSV transmission potential R_0_, the available proportion of resistant individuals and their distribution across herds. PRRS prevalence is defined as the proportion of herds with effective disease transmission potential R above 1. The four graphs show four different distribution scenarios of resistant animals into herds (see Table 1 and text for details). a) Optimal distribution, b) Comprehensive distribution, c) Concentrated distribution, d) Unregulated distribution. Shaded areas correspond to confidence intervals comprising 95% of the predicted values from 100 simulated replicates (note that in a to c, these are too narrow to be visible). Note that in the unregulated distribution scenario (Fig. 1d), the actual proportion of genetically resistant animals across all herds may be lower than the available proportion (presented on the x-axis), explaining why elimination is not possible even if there is unlimited supply of genetically resistant pigs.

Specifically, under optimal conditions where the herd-specific *R*_*0*_ is known or accurately estimated, resistant animals, if sufficiently available, can be distributed according to demand to reduce the herd-specific *R* to below one (see above equation, or eq [1] in Methods). This *optimal distribution* leads to a significant reduction in disease prevalence even under higher average *R*_*0*_ (Figure 1a), and could achieve disease elimination when less than half of the national pig population is genetically resistant for relatively high average *R*_*0*_ (i.e. avg. R_0_ <3). For a moderate average *R*_*0*_ of 1.5, as estimated for PRRS (28, 29), the required proportion of genetically resistant pigs drops to 30% (Figure 1a). While this ideal situation would provide the best environment for PRRS elimination using genetically resistant animals, it is unlikely to occur in a real pig production system, where the herd specific *R*_*0*_ is unknown and farmers can be expected to differ in their willingness and capability to invest in adopting the new technology. A perhaps more realistic distribution scenario, hereafter called *comprehensive distribution*, assumes that all herds are supplied with an equal proportion of available genetically resistant animals, and the sourcing of resistant pigs is managed by the supplying breeding companies rather than the farmer (Figure 1b). Under these circumstances, disease prevalence only decreases considerably when the available proportion of genetically resistant individuals in the population is high. In particular, disease elimination is only possible if the majority of individuals are genetically resistant (e.g. 74% for average *R*_*0*_ of 1.5 (Figure 1a)). The third alternative model scenario, hereafter called *concentrated distribution*, considers that not all farmers may embrace gene-editing and splits farmers into “adopters” and “non-adopters”. Randomly chosen adopters are supplied with an equal fixed proportion of genetically resistant animals (where the proportion may or may not be based on national or regional estimates for *R*_*0*_), whereas non-adopters opt out of this supply entirely. In contrast to the other scenarios, this concentrated distribution leads to a linear reduction in disease prevalence with increasing proportion of genetically resistant animals. Disease elimination is however unachievable unless the supply is based on reasonably accurate estimates of *R*_*0*_. For moderate average *R*_*0*_ of 1.5 this implies that most herds (>98%) would need to contain a large proportion (∼75%) of genetically resistant animals (Figure 1c & Figure 2a). In contrast to all regulated scenarios, the fourth scenario simulated an entirely *unregulated distribution* of genetically resistant animals, where adoption of these animals was assumed to be entirely optional to the farmer. Thus, from a modelling perspective, arbitrary herds are supplied with arbitrary proportions of resistant animals independent of herd size or herd-specific *R*_*0*_. This scenario leads to a relatively small reduction in disease prevalence with high uncertainty, as represented by the wide confidence intervals in the simulations (Figure 1d). Disease elimination through gene editing alone is out of reach for this unregulated distribution scenario.

**Figure 2.**
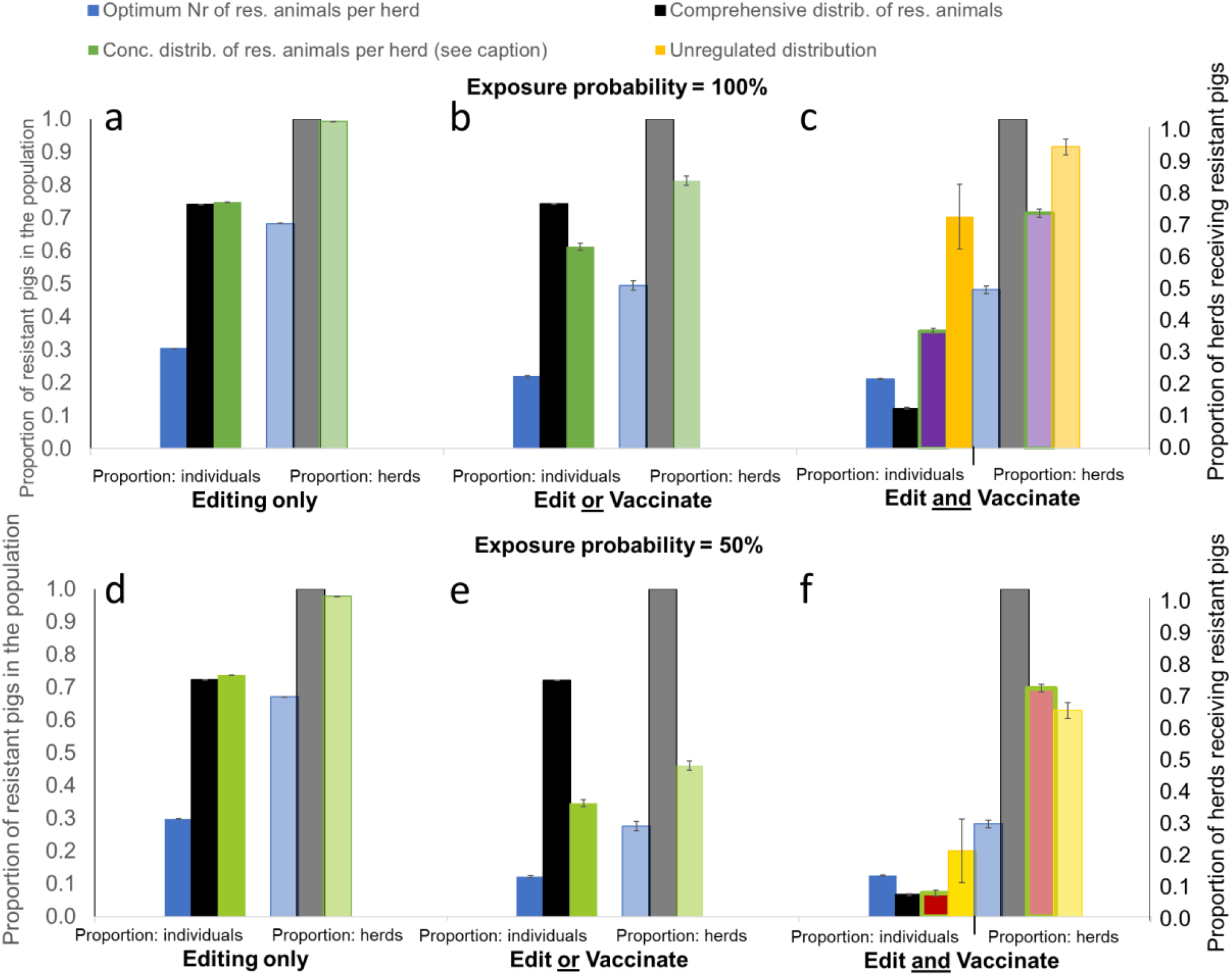
Minimum required proportion of genetically resistant animals (solid bars) and corresponding herds adopting gene editing (transparent bars) for achieving disease elimination through gene editing alone or with vaccination combined, depending on how edited animals are distributed across the herds. Results are shown for average R_0_ value of 1.5 and exposure probability of either 100% (Fig.2 a-c) and 50% (Fig.2 d-f), and vaccine effectiveness of 70%. Different colours refer to different distribution scenarios (see Table 1) with blue = Optimum, black = Comprehensive, green = Concentrated and yellow = Unregulated. The proportion of edited animals in the Concentrated scenarios is chosen at the smallest possible proportion for elimination under each scenario, resulting in a P_e_ of 0.75 for scenarios a, b, d, e (green bars), a P_e_ of 0.5 for scenario c (green bars, purple fill) and a P_e_ of 0.1 for scenario f (green bars, red fill). For further explanation of editing and vaccination strategies, and the different distribution of edited individuals across herds see text.

The above model scenarios assume a pessimistic situation where all herds are exposed to *PRRSV* infected pigs. Reducing the exposure probability of each herd to 50% had however little effect on the overall model predictions: unless the baseline transmission potential *R*_*0*_ is known and the distribution of genetically resistant pigs is regulated accordingly, PRRS elimination through gene editing alone is only achievable if almost all herds (>95% for *R*_*0*_ = 1.5) consist primarily (>70% for *R*_*0*_ = 1.5) of genetically resistant animals (see Figure 2d-f for moderate *R*_*0*_ = 1.5 and Fig. S1 for high *R*_*0*_ = 5).

### Gene editing and vaccination as combined disease control

Controversial technologies such as gene editing are unlikely to fully replace existing measures of disease control soon. The second question we therefore sought to answer is how gene editing can effectively complement existing disease control measures. Mass vaccination of pigs against PRRS is already widespread in many countries, but has limited effectiveness (Rowland & Morrison, 2012) and subsequently cannot serve as a singular elimination tool. To investigate the combined impact of gene editing and vaccination on the feasibility of eliminating PRRS, we calculated the *PRRSV* transmission potential (see Methods) for scenarios where either vaccination or gene editing are applied as the sole disease control strategies or applied either as complementary alternatives (hereafter referred to as *Edit or Vaccinate* scenario, see Methods), or jointly (hereafter referred to as *Edit and Vaccinate* scenario, see Methods).

In line with existing estimates, our model (with *R*-values calculated using equation [1] in Methods) predicts that PRRS elimination cannot be achieved through mass vaccination alone when vaccine effectiveness is 70% or less and the average *R*_*0*_ is 1.5 and exposure probability is 50% or higher (31). Elimination is however achievable if vaccination and gene editing are deployed together (Figure 2). Compared to the requirements for eliminating PRRS through gene editing alone, the required amount of genetically resistant animals and herds adopting such animals reduces considerably if gene editing is complemented by mass vaccination (Figure 2). The biggest gains occur if vaccination is applied to all susceptible animals (*Edit and Vaccinate* scenario, Figure 2c &f) rather than just in herds that deploy vaccination as an alternative disease control to gene editing (*Edit or Vaccinate* scenario, Figure 2b & e). For example, when the average *R*_*0*_ is 1.5 and all herds are exposed to *PRRSV* infection, the required proportion of genetically resistant pigs drops by 83% from 74% to as little as 12% resistant pigs for the centrally regulated *Comprehensive* distribution scenario when gene editing is complemented by vaccination of all susceptible animals with a vaccine of 70% effectiveness (Figure 2c).

Perhaps most importantly, the model predicts that PRRS elimination becomes possible even when the adoption of genetically resistant animals is unregulated if mass vaccination is simultaneously applied, although it would still require most herds in a population to purchase genetically resistant animals (Figure 2 c&f). The exact percentage of herds and genetically resistant animals required depends strongly on the baseline transmission potential (See Figure 2 and Figure S1) and the exposure probability. Whereas the voluntary scheme would require 70% of pigs to be genetically resistant in over 91% of herds when the average *R*_*0*_ is 1.5 and *PRRSV* exposure is 100%, only 20% of genetically resistant pigs distributed across 63% of herds would suffice if the exposure probability dropped to 50% (Fig. 2c&f).

As would be expected, the required number of resistant pigs increases when the transmission potential of PRRS is higher. However, even in a severe scenario corresponding to average *R*_*0*_ of 5 and 100% exposure, the model predicts that disease can be eliminated when all herds are supplied with a set proportion of 53% genetically resistant animals and all susceptible pigs are vaccinated (see Supplement Figure S1).

### Impact of vaccine effectiveness on disease elimination

Whereas gene editing and vaccination with vaccines of relatively high effectiveness (≥70%) emerges as a highly effective PRRS elimination strategy in our models, vaccination with poorly effective vaccines is predicted to contribute relatively little to disease elimination. This is illustrated in Figure 3 (and Figure S2 for higher R_0_), which also shows that for a voluntary distribution scheme, disease elimination is no longer possible when vaccine effectiveness is 50% or less.

**Figure 3.**
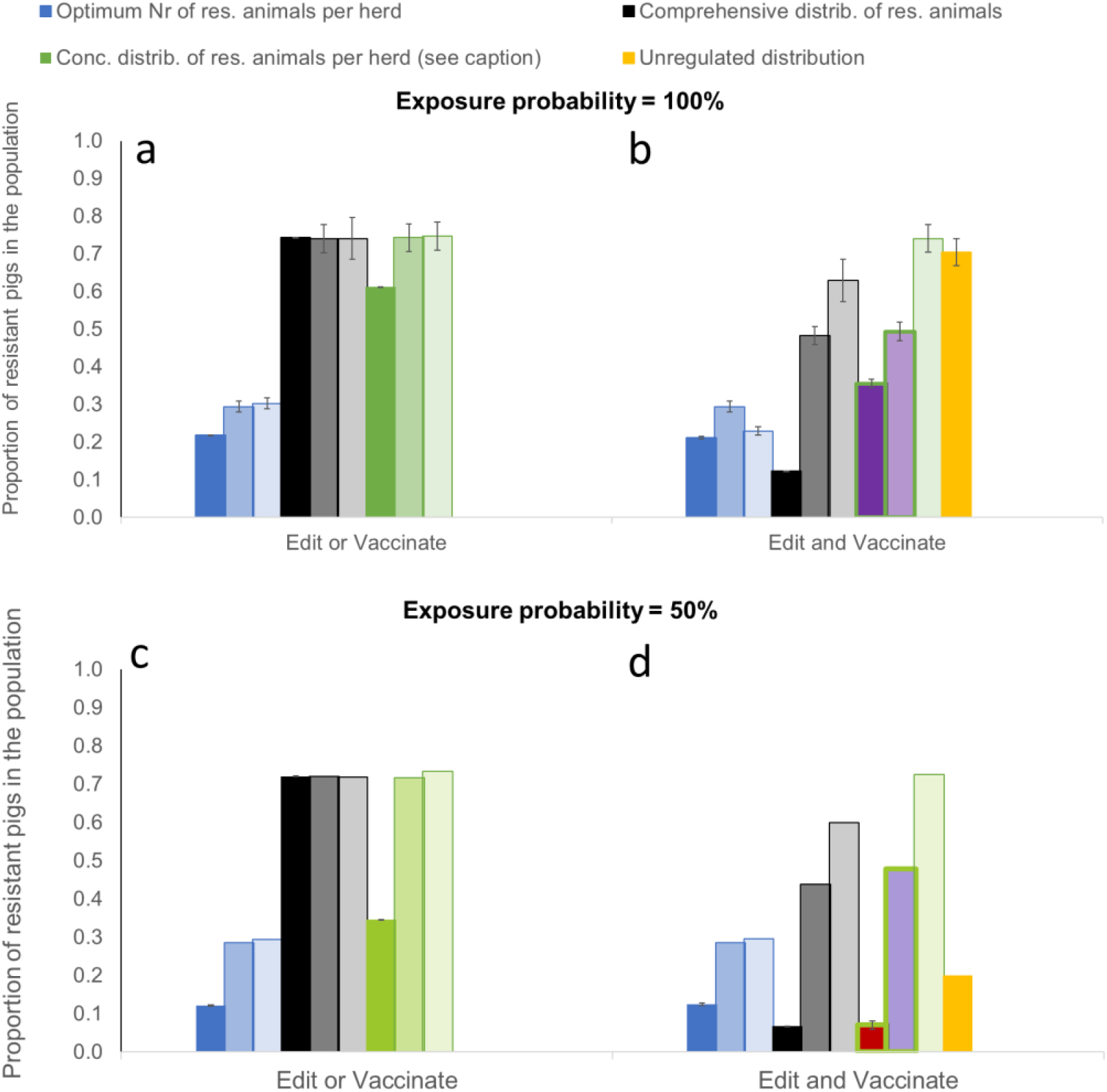
Minimum required proportion of genetically resistant animals for achieving disease elimination through gene editing and vaccination combined, depending on vaccine effectiveness ε_V_ and exposure probability. Dark bars: ε_V_ = 0.7, medium bars: ε_V_ = 0.5; light bars: ε_V_ = 0.3. Different colours refer to different distribution scenarios (see Table 1) with blue = Optimum, black = Comprehensive, green = Concentrated and yellow = Unregulated. The proportion of edited animals in the Concentrated scenarios is chosen at the smallest possible proportion for elimination under each scenario, resulting in a P_e_ of 0.75 for scenarios a, c (green bars), a P_e_ of 0.5 for scenario b (green bars, purple fill) and a P_e_ of 0.1 for scenario d (green bars, red fill). An average transmission potential of R_0_ = 1.5 was assumed.

### Time scale for achieving disease elimination

With the required proportions of genetically resistant animals under different strategies defined, the third question surrounding the feasibility of gene editing can be addressed: How long does it take to produce the required numbers of genetically resistant animals using current breeding techniques? Given the potentially limited shelf life of gene editing caused by the emergence of escape mutants, fast dissemination of genetically resistant pigs into the commercial level is crucial. This could be hampered by the fact that gene editing technology will be limited to the top tier of the multi-tier pig production pyramid (Figure 4) and that the PRRS resistance allele is recessive (14). Genotyping of pigs to trace resistance genotypes could help to identify both resistant and heterozygous carrier selection candidates and propagate the resistance allele efficiently through the production pyramid. However, genotyping is costly and not usually applied in the lower tiers. Despite these limitations, which were considered in our gene flow simulation model (see Methods for details), we found that gene edited resistance alleles can be efficiently disseminated through the tiers of the population without continuous genotyping of selection candidates in lower population tiers. Through selective mating of both homozygous resistant and heterozygous carrier animals in the top two tiers where genotyping is conventionally carried out, the resistance allele effectively propagates through the whole production pyramid, eventually resulting in genetically resistant animals carrying two copies of the resistance allele in the commercial tier (Figure 4).

**Figure 4.**
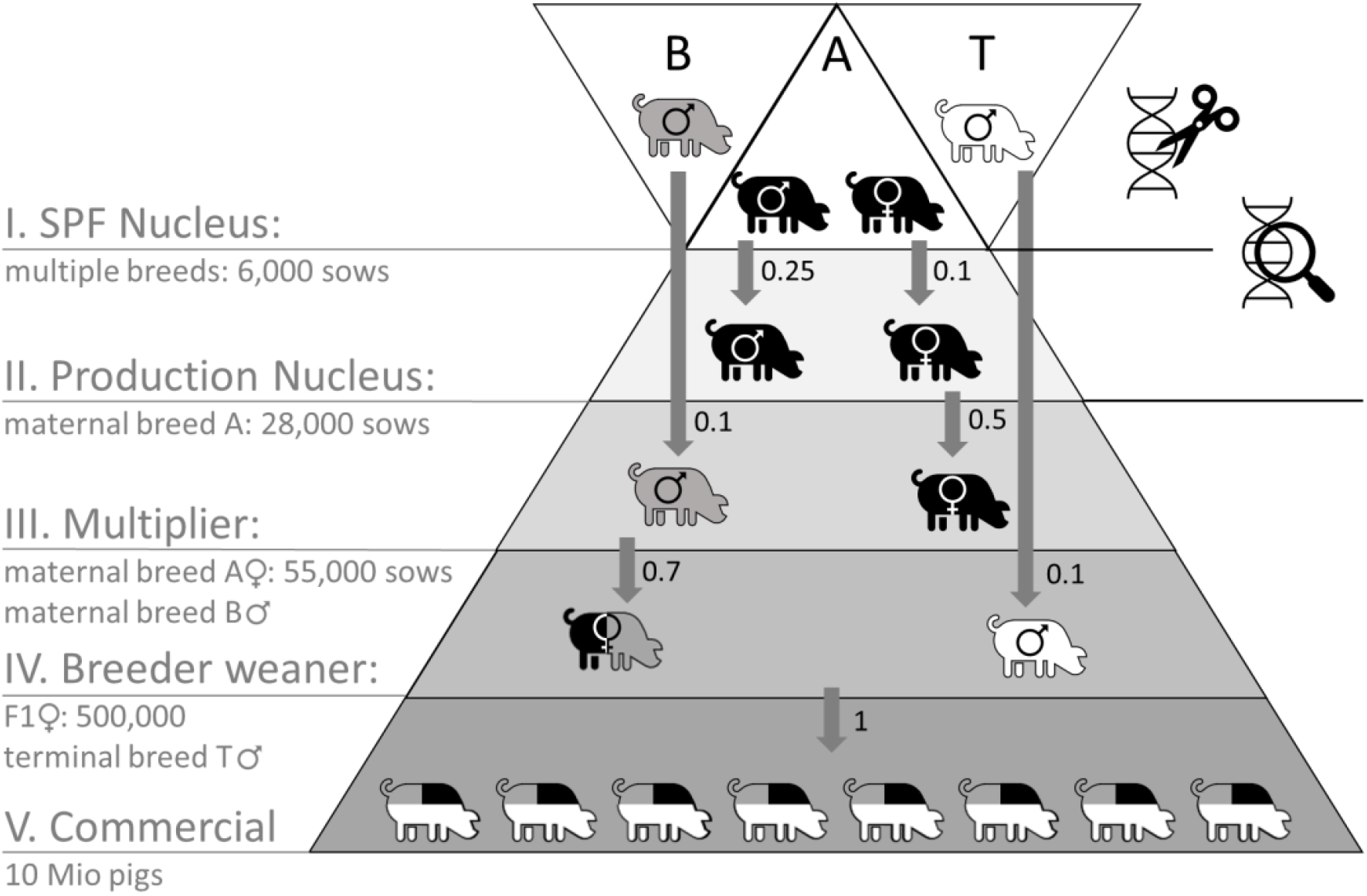
Schematic diagram of a typical five-tier pig production structure implemented into the gene flow model. Two breeds, A and B, are crossed to create hybrid females that are mated to males from a third breed T to produce commercial animals. Numbers next to the arrows denote the selection proportions transferred into subsequent tiers. Gene editing is limited to tier I only; genotyping of selection candidates is carried out in tiers I and II.

Our natural gene flow model predicts that the required proportion of resistant animals in the commercial population to achieve disease elimination under the different distribution and vaccination scenarios above can be reached within less than 6 years (see Figure 5). In the best case scenario, where genetically resistant animals are distributed optimally across herds and this is augmented by mass vaccination with a vaccine of at least 70% effectiveness (either only in herds that do not receive resistant animals or of susceptible animals in general), this can be achieved within less than 3 years (green lines, Figure 5, for details on timepoints see Table S1 in the supplement). Gene editing a higher percentage of selection candidates in the top tier of the production pyramid does not result in a proportional reduction of the time needed to produce the required proportion of resistant animals (e.g. in the example above, increasing the editing proportion from 5% to 20% only reduces the time before required numbers are reached by 20%).

**Figure 5.**
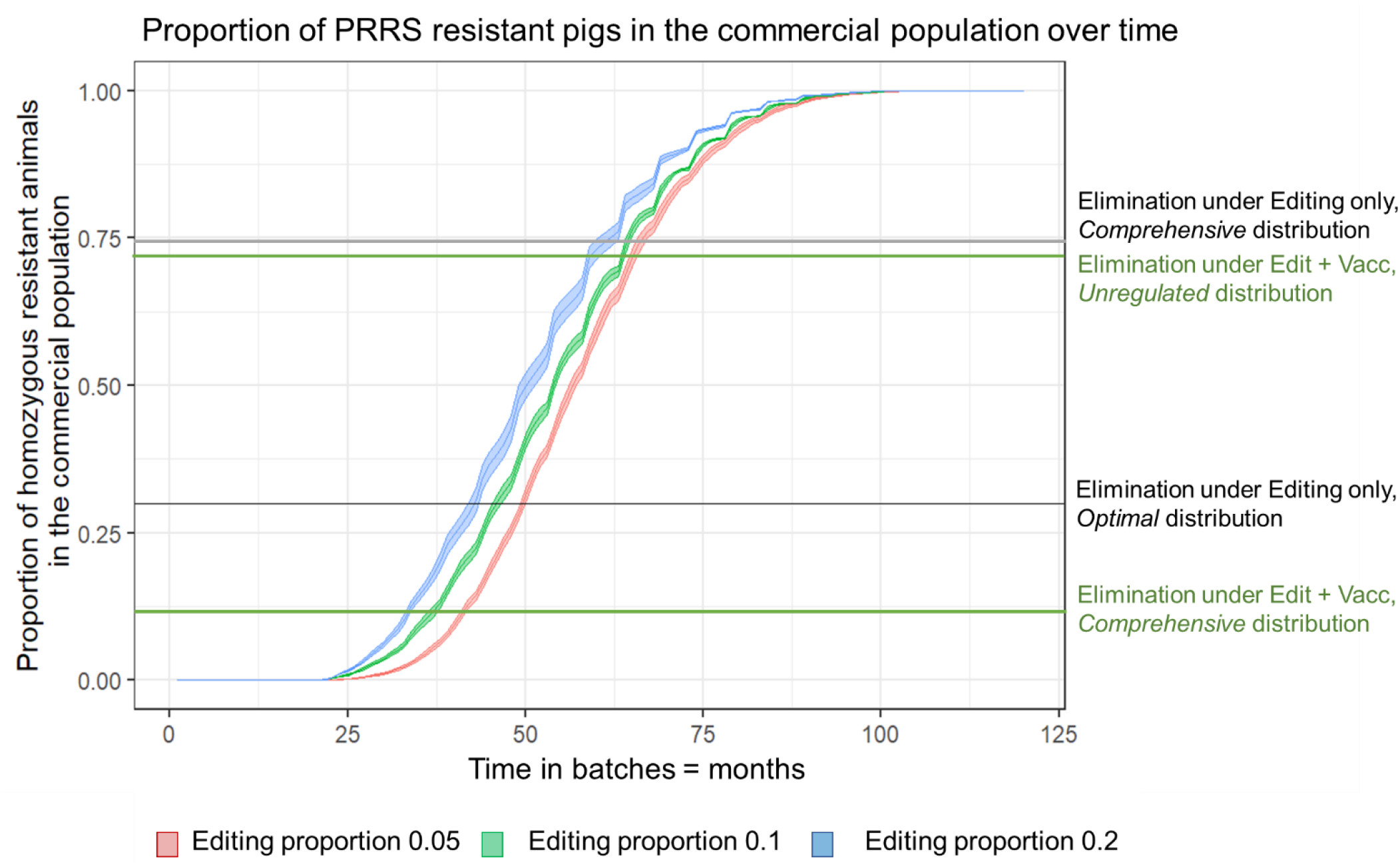
Time to reach proportions of resistant pigs in the population needed for PRRS elimination under different gene editing scenarios. The indicated thresholds levels refer to required numbers of genetically resistant pigs for achieving elimination under different distribution scenarios of pigs in the commercial tier (average R_0_ = 1.5 and exposure probability = 100%). For visibility, not all scenarios are depicted.

## Discussion

The results of our modelling study suggest that gene editing could drastically reduce PRRS prevalence and may succeed in eliminating PRRS within three to six years of selective breeding. If gene editing was the only disease elimination tool, this would however require a highly regulated distribution scheme that supplies the majority of herds with a disproportionally large percentage of genetically resistant pigs. Given that adoption by farmers remains one of the biggest barriers to implementation of biotechnology (32), this blanket distribution of a novel genomic technology seems unlikely under current conditions. Nonetheless, we found PRRS elimination still to be feasible for a more realistic scenario where gene editing and mass vaccination are used conjunctively, allowing individual farmers to choose their management tool. Effective application of both control strategies simultaneously drastically reduces the required number of genetically resistant pigs and herds needed to adopt these and can achieve elimination even without stringent regulations concerning their distribution. Since PRRS has proven difficult to combat with conventional disease control (19, 33), this finding is encouraging, as it illustrates that effective combination of existing control tools with novel genomic technologies may achieve the so far impossible outcome of much desired disease elimination.

Our model, despite its simplicity, provides important first insights into the key determinants and their interactions that underpin the success of gene editing in controlling livestock epidemics.

### Determinant 1: The baseline transmission potential R_0_

As expected, the higher the baseline transmission potential *R*_*0*_, the more stringent control measures (e.g. more genetically resistant pigs) are needed to achieve a desired outcome (compare Figure 2 (mean *R*_*0*_ = 1.5) and Figure S1 (mean *R*_*0*_ = 5)). In practice, the implementation of effective disease control is hampered by the fact that *R*_*0*_ typically varies across sub-populations and that precise estimates of *R*_*0*_ are rarely available (34, 35). Our model accommodates for heterogeneities in *R*_*0*_ implicitly by drawing herd-specific *R*_*0*_ values from normal distributions. The results highlight the importance for obtaining precise sub-population specific estimates of *R*_*0*_, as such estimates allow for more effective targeted disease control with minimum wastage of valuable resources, such as genetically resistant pigs. The *Optimal* distribution scenario in our model, which assumes that herd specific *R*_*0*_ values are known, required up to 60% fewer genetically resistant pigs for disease elimination compared to other distribution scenarios with less precise or no knowledge of *R*_*0*_. However, given the high uncertainty in herd-specific *R*_*0*_-values in practice (35), we incorporated different degrees of knowledge about *R*_*0*_ in the modelled distribution scenarios, ranging from full knowledge of herd specific *R*_*0*_ represented by the *Optimum* distribution scenario to partial knowledge (e.g. national or regional average *R*_*0*_ estimates) accommodated within the *Concentrated* scenario to potentially zero knowledge represented by the other scenarios. Based on our model predictions, PRRS elimination through gene editing was only possible if *R*_*0*_ was at least partially known or complemented by mass vaccination of all susceptible individuals with a sufficiently effective vaccine.

### Determinant 2: Distribution of genetically resistant animals across herds

Our model results show that reduction in disease prevalence and the prospect of disease elimination depend strongly on how available genetically resistant animals are distributed across herds. Whereas the modelled *Optimum* distribution was able to eliminate PRRS from a national commercial pig population without complementary vaccination with only as little as 30% of pigs carrying the PRRS resistance genotype, the *Unregulated* distribution could only achieve elimination if 70% of all pigs were genetically resistant and the remaining pigs were vaccinated. Feasibility issues with regards to the appropriate dissemination of genetically resistant individuals in commercial populations warranted modelling a variety of potential scenarios.

The *Optimum* distribution scenario provides valuable insights into the potential scope of gene editing for controlling epidemics in a hypothetical world, where the full-scale benefits of gene editing for disease control can be realized. However, it is unlikely to be met in practice as it not only assumes that herd specific *R*_*0*_ values are known, but also that PRRS resistant pigs are identifiable, and that no obstacles for providing each herd with the required number of genetically resistant pigs exist. Identifying PRRS resistant pigs would require either tracing the parentage across the production pyramid or genotyping all commercial pigs, neither of which are current industry practices. Unless adoption of genetically resistant pigs was made compulsory (*Comprehensive* scenario), only a fraction of herds is therefore likely to contain these pigs in practice. Furthermore, the proportion of genetically resistant pigs that each of these herds receive could be either controlled by the supplier (*Concentrated* scenario) or by the farmer (see *Unregulated* scenario). Either of them could base their decisions on estimates of *R*_*0*_, which are realistically only available on a national or regional level. Our choice of distribution scenarios aimed to capture this wide spectrum of potential scenarios, and to provide useful quantitative estimates of the associated impact. To accommodate the common lack of herd specific *R*_*0*_ estimates, all distribution scenarios except for the *Optimum* scenario assumed that the proportion of genetically resistant pigs per herd is independent of the herd specific *R*_*0*_. It should be noted that predictions for all alternative scenarios to the *Optimum* scenario also apply if the resistance genotype of pigs was not exactly known, as long as the overall proportion of genetically resistant pigs in the population was known by the suppliers and pigs were distributed randomly to the receiving herds. Our model provides quantitative estimates how each distribution scenario may affect PRRS prevalence and importantly reveals that gene editing can substantially reduce the prevalence even if adopted in restricted, sub-optimal capacity.

### Determinant 3: Combination of alternative control measures with different effectiveness

There is general acceptance that no single silver bullet can eliminate persistent diseases such as PRRS, but that this would require a combination of effective control measures (36–38). Correspondingly, our epidemiological model predicts that PRRS elimination cannot be realistically achieved through the sole application of gene editing or vaccination but becomes feasible if both measures are effectively used in conjunction. Importantly, our results suggest that the likely presence of staunch non-adopters, e.g. farmers that cannot be incentivized to participate in an elimination scheme on the basis of gene editing, may not necessarily stand in the way of realising the full-potential of gene editing since not all herds have to receive genetically resistant animals if simultaneous vaccination is applied.

Our model results also demonstrate that the success of combined control strategies hinges on their relative effectiveness. Whereas evidence to date suggests that pigs carrying two copies of the PRRS resistance alleles are fully resistant to *PRRSV* infection (effectiveness of gene editing is one) (23, 24, 30), the effectiveness of existing PRRS vaccines is compromised amongst other factors by the limited cross-protectivity of a given vaccine to different *PRRSV* strains (39, 40). In our model, elimination of PRRS falls out of reach for the less stringent *Unregulated* and *Concentrated* distribution scenarios if vaccine effectiveness drops below 50%. These predictions clearly demonstrate the need for continued support of vaccine development even when new and perhaps at first sight more promising technologies such as gene editing appear on the horizon.

Similar to gene editing, the impact of vaccination also depends strongly on vaccine coverage (Bitsouni et al., 2019). Here we deliberately made the strong assumption that mass vaccination is applied either in all herds that don’t adopt gene editing, or in all herds altogether. Although PRRS vaccination is wide-spread in practice, these assumptions are obviously unlikely to be met in reality. Incomplete vaccine coverage would prevent disease elimination when the adoption of genetically resistant pigs is sparse and exposure risk is high (results not shown). This highlights the need to consider additional determinants that may underpin the success of gene editing for disease control in future studies, such as natural genetic variation in pigs’ PRRS resistance. Indeed, genetic selection of pigs for increased natural PRRS resistance has been advocated as a viable complement to existing PRRS control (41, 42). Combined application of these complementary genetic disease control strategies may effectively eliminate PRRS even under restricted vaccine usage.

### Determinant 4: Exposure risk

It is unlikely that all herds are simultaneously and equally exposed to the PRRS virus. Heterogeneity in exposure was included in our model through a random uniform exposure probability distribution. Whilst reduction of the average exposure risk from 100% to 50% had little influence on the model results associated with gene editing as sole disease control strategy, it drastically reduced the requirements for genetically resistant animals when gene editing and vaccination were used in conjunction. In reality, exposure risk will likely depend on PRRS prevalence in herds that are in close spatial proximity or linked through e.g. transport or trading (43, 44). Whilst spatial factors were not explicitly considered in our model presented here, exploration of these is an important avenue for future modelling studies as they would allow more strategic and targeted distribution of genetically resistant animals in epidemic hotspots.

### Timeliness and other considerations

*PRRSV* has been estimated to have the highest evolutionary rate (on the order of 10^−2^/site/year) of all known RNA viruses (with rates ranging from 10^−3^ to 10^−5^/site/year) (45). This alarming evolutionary rate, together with observations that the virus evolves towards increased virulence with ability to evade vaccine-induced immunity (46), raises concerns about how long the current gene editing process confers complete resistance to this virus. Hence, ambitious goals such as disease elimination, would need to be achievable within a short time frame. Coupling the epidemiological model with a gene flow model suggests that PRRS can be potentially eliminated through use of gene editing within three to six years. Although impossible to predict whether this is sufficiently fast to win the race against virus evolution, this time scale fits well within the anticipated time scale of current national or regional elimination programmes for PRRS and other livestock diseases (38, 47).

A number of simplifying assumptions in our gene flow model warrant further investigations with regards to their impact on the predicted time scales. Our model describes the national pig industry by a five-tier breeding pyramid originating from three pure breeds. Although this structure is common for modelling pig breeding schemes (48, 49), it does not take into account the multitude of different breeding companies and different breeds that often form part of the crossbreeding schemes behind hybrid pig production. Furthermore, we assumed that all selection candidates for selection in the top pyramid tier are also candidates for gene editing, thus ignoring the possibility that some breeding companies may not apply the technology to all selection candidates, or not apply it at all if this meets best their costumers’ demand. Our model could easily accommodate this increased complexity by increasing the proportion of gene edits carried out to a subset of selection candidates in the top tier. In the current model gene editing of 20% of animals in the top tier was sufficient to satisfy the demands for genetically resistant animals in the lower tiers. Increasing this proportion in a subset of breeds composing the top production tier would generate the required number of genetically resistant animals in the commercial population in a similar time frame.

Our gene flow model assumes gene editing technologies to be incorporated into traditional breeding schemes based on natural mating or artificial insemination of selection candidates. However, a number of more efficient methods for fast propagation of genetically resistant to the commercial tier have been recently proposed, such as e.g. the use of surrogate sire technology (50) or gene-drives (51) for the faster propagation of the resistance allele, or the use of e.g. self-terminating “daisy chain” gene-drives that disappear from the population after a few generations (52). These may not only accelerate the rate at which genetically resistant animals can be produced, but may also help to limit potential contamination effects of gene editing on the wider population (53), e.g. organic producers that need to ensure that their animals do not carry any artificially altered genetic material.

Lastly, it is important to remind readers that this study focused purely on the epidemiological impact of gene editing. Implementation of this highly controversial technology into practical disease control will also largely depend on economic and societal aspects. Future studies are therefore required to assess the economic feasibility of the approaches presented here and to weigh the associated economic costs against the benefits of eliminating one of the costliest livestock diseases to date. One major cost factor flagged up by our models concerns the investment into routine genotyping of commercial pigs, which would allow identification and targeted distribution of genetically resistant pigs, thus increasing the chance of disease elimination. In addition, future studies should consider potential trade-offs arising from gene-editing with genetic improvement in other important livestock traits. Preferential selection of animals carrying the resistance allele likely results in a loss in genetic gain for other traits in the breeding goal. The scale of this trade-off will strongly influence the willingness of livestock breeders to produce genetically resistant animals.

## Conclusions

In summary, our proof-of-concept study highlights hitherto unprecedented opportunities for eliminating infectious diseases in livestock by complementing existing control methods with novel gene editing technologies. The model provides some first quantitative estimates of how many edited individuals may be required, and how these would need to be distributed depending on the overall transmission potential of the disease and the quality and application of available vaccines. It particularly highlights the continued need to develop vaccines with high effectiveness, and to consider additional control options such as genomic selection for natural (yet incomplete) PRRS resistance. Effective combination of these alternatives increases the chance for disease elimination and reduces the requirements for stringent regulations concerning the application of each of these measures. Finally, our study provides some first estimates of resource requirements to balance epidemiological benefits against economic trade-offs and stresses the urgent need to carefully investigate and weigh epidemiological and economic benefits against ethical and other societal concerns.

## Methods

### The epidemiological simulation model

We simulated a commercial pig population representative for many countries in Europe or pig-producing regions in North America or China (54), which consisted of 12 million pigs distributed into 5,000 herds. Herd size was assumed normally distributed around a mean of 2,400 with a standard deviation of 1,000 pigs. Furthermore, we assumed that each herd is exposed to *PRRSV* infection with a given exposure probability *p*_*exp*_. This value was originally set to one to model the worst-case scenario and then reduced to 0.5 to mimic the more realistic situation of heterogenous exposure risk.

Once exposed, epidemiological theory stipulates that an infectious disease cannot invade a herd if its transmission potential, i.e. the so-called reproductive ratio *R*, is below one, whereas invasion is possible when *R* > 1 (27). The *R*-value is usually not precisely known and is expected to differ between individual herds, depending on the circulating pathogen strain, the pig breed, individual variation in resistance to the infection, environmental factors, as well as herd management and biosecurity characteristics (31, 55). Detailed epidemiological modelling of *PRRSV* transmission dynamics that considers these demographic characteristics as well as within- and between herd contact structures affecting disease transmission will be an important avenue for future predictive modelling, but as a first step we here sought to gain initial qualitative and quantitative understanding about the potential impact of gene editing on PRRS control. To achieve this, we simply assumed that in the absence of gene editing or vaccination, the baseline *PRRSV* transmission potential *R*_*0*_ for the different herds follows a normal distribution ∼N(μ_R0_, σ_R0_), which is independent of the herd size, i.e. PRRS transmission was assumed to be density-dependent (56).

Following epidemiological theory (27) and assuming no interactive effects between genetic resistance and vaccination, the presence of genetically resistant and / or vaccinated pigs in a herd reduces the herd specific R_0-_value to the effective reproductive rate

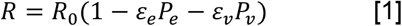

where the parameters *ε*_*e*_ and *ε*_*v*_ denote the effectiveness (i.e. proportional reduction in *PRRSV* infection) of gene editing and vaccination, respectively, and *P*_*e*_ and *P*_*v*_ denote the fraction of genetically resistant or vaccinated animals in the current herd, respectively, with *P*_*e*_ + *P*_*v*_ ≤ 1. For scenarios representing heterogeneous exposure, herds (chosen at random with probability *p*_*exp*_) that are not exposed to *PRRSV* infection are assigned a value *R* < 1. Input parameters with the assumed ranges for the epidemiological simulation model are listed in Table S2.

In this study we define PRRS prevalence as the proportion of herds for which the effective reproductive rate *R* ≥ 1 as per equation [1]. *PRRSV* is considered to be eliminated from the population if *R* < 1 in over 99% of herds.

Equation [1] allows calculation of the required proportion of genetically resistant and vaccinated individuals to achieve herd immunity, i.e. *R* < 1. In particular, in a non-vaccinated herd (*P*_*v*_ = 0) and assuming gene-editing efficacy *ε*_*e*_=1, the required minimum proportion of edited pigs per herd for preventing disease invasion (i.e. achieving *R* < 1) is

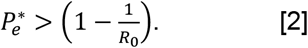

Expression [2] implies that PRRS can in principle be eliminated from a national pig population if the herd specific R_0_-values were known or could be reliably estimated and each herd contains the critical number of genetically resistant individuals 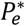.

According to [1] and [2], disease prevalence and elimination on a national scale depend not only on the proportion of genetically resistant and vaccinated animals in a population, and on the effectiveness of the corresponding control measure, but also on how these animals are distributed across the herds. The proportions *P*_*e*_ of genetically resistant pigs in each herd were specified by the corresponding distribution and vaccination scenarios. Specifically, for the *Optimal, Concentrated* and *Unregulated* distribution scenarios (Table 1), herds were selected at random to receive the required proportion 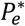 (*Optimal*), or a given fixed proportion *P*_*e*_ (*Concentrated)*, or arbitrary proportion *P*_*e*_ (*Unregulated)* of genetically resistant animals, respectively, until the available stock of genetically resistant animals was either depleted or the demand was satisfied, whichever fast achieved first. In the *Comprehensive* distribution strategy, the available stock of genetically resistant animals was distributed uniformly across all herds thus resulting in an average equal fraction of edited animals *P*_*e*_ in each herd.

For simulations that also included vaccination, the distribution of genetically resistant animals across herds was carried out first, and vaccination was subsequently assumed to be applied to either all animals in herds that contained no genetically resistant animals (*Edit or Vaccinate* strategy) or to all remaining susceptible animals across all herds (*Edit and Vaccinate* strategy). Thus, for the *Edit or Vaccinate* strategy the proportion *P*_*v*_ of vaccinated individual per herd is either zero or one, depending on whether the farmer adopts genetically resistant animals or applies mass vaccination to control PRRS. For the *Edit or Vaccinate* scenario, where all non-resistant animals (and possibly also resistant animals if their resistance status is unknown) are vaccinated, *P*_*v*_ was set to 1 − *P*_*e*_.

For each model scenario, 100 replicates were produced, and the means and standard errors over the replicates were calculated. A differential genetic algorithm was used (57) to calculate the minimum number of herds and genetically resistant animals required to achieve disease elimination for each simulated scenario.

### Gene flow simulation model

We developed a stochastic gene flow simulation model to track the propagation of PRRS resistance alleles through a typical 5-tier pig production pyramid into the commercial pig population (48), where gene editing can realistically only be carried out on a subset of pigs at the top pyramid tier. This specific pathogen-free (SPF) nucleus tier typically consists of purebred animals (here from three distinct breeds) that are selectively bred at high health and management level, and for which a proportion are then sold or provide semen to farms in the lower tiers of the pyramid, as outlined in Figure 4. Pigs in each tier are produced through mating (or artificial insemination of) a fixed proportion of males and females from the same or upper pyramid tiers that have been selected to act as parents for the next generation (see Table S3 for selected proportions and mating ratios), thus propagating their genetic material to offspring in the same or subsequent tier.

To assign a timescale to the natural propagation of the resistance alleles through the production pyramid, offspring in each tier are produced in the simulations in discrete monthly batches to represent a management system that is aligned with the natural reproductive and life cycle of pigs (see assumed parameter values in Table S4).

PRRS resistance was assumed to be controlled by a single gene in our model and to follow Mendelian inheritance patterns. Since PRRS resistance is expected to be just one of multiple genetic traits on which selection decisions are based, each animal was also assigned a single value representing its total genetic merit that it passes on to its offspring. This value represents a combination of genetically correlated and uncorrelated traits controlled by many genes with standard polygenic inheritance patterns (58) and allows for the calculation of mean genetic merit of the entire population.

In the beginning of the simulation, a stable starting population was generated for each tier of the production pyramid in the absence of gene editing. This was achieved by first creating founder populations for each of the three breeds represented in the top pyramid tier, where each animal was assigned a genetic merit drawn from a random normal distribution. Specified proportions of individuals were then selected for mating within the nucleus based on their genetic merit (for proportions, see Table S3). Once a stable base population was obtained within the nucleus (after about 18 months), individuals (or semen) were selected for transfer to subsequent tiers as shown in Figure 4. The burn-in phase was then run for an additional 33 months to create base populations in all pyramid tiers. The maximum numbers of sows in each tier were back-calculated based on the number of commercial piglets produced annually (12 Million), the selection proportions, and the underlying pig life cycle parameters (Table S4). The burn-in period resulted in a homogenously susceptible population that contained no animals carrying the PRRS resistance allele.

PRRS resistance was introduced into the population by selecting a fixed proportion (5%, 10% or 20%, respectively) of animals with the highest genetic merit from each breed in the top tier of the production pyramid, the SPF Nucleus, to undergo the gene editing process. Gene editing was limited to tier I to test the feasibility of reaching sufficient numbers of resistant animals in the commercial tier without applying repeated gene editing throughout the breeding pyramid. Editing success using CRISPR/Cas9 and embryo survival rates were assumed to be 0.81 and 0.61, respectively (59). Animals in the top pyramid tier were then preferentially selected based on their PRRS resistance genotype as well as (if there were not enough animals carrying at least one PRRS resistance allele) their genetic merit value, thus allowing the resistance alleles to be naturally propagated to the subsequent tiers following Mendelian inheritance patterns.

As selection candidates in tier II, the Production Nucleus tier, were assumed to be genotyped to determine their resistance genotype, preferential selection for the successful gene edit occurred here as well. Since only high-merit selection candidates are selected in the gene editing process inside the SPF nucleus, selection in the absence of genotyping in the lower tiers of the pyramid is expected to also be skewed towards animals carrying the PRRS resistance allele. However, genotyping in the top 2 tiers accelerates the flow of resistant individuals from the top of the breeding pyramid into the lower tiers while reflecting current industry practices.

In tiers III and IV, animals were selected based on their genetic merit alone. The gene flow simulation model generated estimates for the number of animals carrying one or two copies of the resistance allele in each pyramid tier, and in particularly for the number of PRRS resistant animals in the commercial population, over time.

## Acknowledgement

The study was funded through the UKRI Industry Strategic Challenge Fund (ISCF) Transforming Food Production Seeding Award. ADW’s and BW’s contribution was funded by the BBSRC Institute Strategic Programme Grant (BBS/E/D/20002172-4 (ISP2)).

## References

1. K. Dhama, et al., Coronavirus Disease 2019–COVID-19. Clin. Microbiol. Rev. 33, 1–48 (2020).

2. L. Xu, et al., CRISPR/Cas9-Mediated CCR5 Ablation in Human Hematopoietic Stem/Progenitor Cells Confers HIV-1 Resistance In Vivo. Mol. Ther. 25, 1782–1789 (2017).

3. S. Jiang, Q. W. Shen, Principles of gene editing techniques and applications in animal husbandry. 3 Biotech 9, 1–9 (2019).

4. A. Dubock, Golden Rice: instructions for use. Agric. Food Secur. 6, 60 (2017).

5. United Nations world food Programme, 2019 - Hunger Map. United Nations world food Program. - Fight. Hunger worldwide. (2019) (August 13, 2019).

6. United Nations, “World Population Prospects 2019:Highlights” (2019).

7. K.. Thomson, World agriculture: towards 2015/2030: an FAO perspective. Land use policy 20, 375 (2003).

8. B. Bahadur, M. V. Rajam, L. Sahijram, K. V. Krishnamurthy, Eds., Plant Biology and Biotechnology (Springer India, 2015).

9. N. de Graeff, K. R. Jongsma, J. Johnston, S. Hartley, A. L. Bredenoord, The ethics of genome editing in non-human animals: a systematic review of reasons reported in the academic literature. Philos. Trans. R. Soc. B Biol. Sci. 374, 20180106 (2019).

10. X. Wang, et al., Multiplex gene editing via CRISPR/Cas9 exhibits desirable muscle hypertrophy without detectable off-target effects in sheep. Sci. Rep. 6, 1–11 (2016).

11. X. Wang, et al., Disruption of FGF5 in cashmere goats using CRISPR/Cas9 results in more secondary hair follicles and longer fibers. PLoS One 11, 1–12 (2016).

12. D. F. Carlson, et al., Production of hornless dairy cattle from genome-edited cell lines. Nat. Biotechnol. 34, 479–481 (2016).

13. C. Proudfoot, C. Burkard, Genome editing for disease resistance in livestock. Emerg. Top. Life Sci. 1, 209–219 (2017).

14. C. Tait-Burkard, et al., Livestock 2.0 - Genome editing for fitter, healthier, and more productive farmed animals. Genome Biol. 19, 1–11 (2018).

15. J. Rushton, The economics of animal health and production, J. Rushton, Ed. (CABI, 2008) https://doi.org/10.1079/9781845931940.0000.

16. D. J. Holtkamp, et al., Economic Impact of Porcine Reproductive and Respiratory Syndrome Virus on U.S. Pork Producers. Anim. Ind. Rep. AS 658 (2012).

17. H. Nathues, et al., Cost of porcine reproductive and respiratory syndrome virus at individual farm level – An economic disease model. Prev. Vet. Med. 142, 16–29 (2017).

18. Z. Guo, X. X. Chen, R. Li, S. Qiao, G. Zhang, The prevalent status and genetic diversity of porcine reproductive and respiratory syndrome virus in China: A molecular epidemiological perspective. Virol. J. 15, 1–14 (2018).

19. R. R. R. Rowland, R. B. Morrison, Challenges and Opportunities for the Control and Elimination of Porcine Reproductive and Respiratory Syndrome Virus. Transbound. Emerg. Dis. 59, 55–59 (2012).

20. T. G. Kimman, L. A. Cornelissen, R. J. Moormann, J. M. J. Rebel, N. Stockhofe-Zurwieden, Challenges for porcine reproductive and respiratory syndrome virus (PRRSV) vaccinology. Vaccine 27, 3704–3718 (2009).

21. J. Christopher-Hennings, et al., Porcine reproductive and respiratory syndrome (PRRS) diagnostics: Interpretation and limitations. J. Swine Heal. Prod. 10, 213–218 (2002).

22. N. J. Boddicker, et al., Genome-wide association and genomic prediction for host response to porcine reproductive and respiratory syndrome virus infection. Genet. Sel. Evol. 46, 18 (2014).

23. C. Burkard, et al., Pigs Lacking the Scavenger Receptor Cysteine-Rich Domain 5 of CD163 Are Resistant to Porcine Reproductive and Respiratory Syndrome Virus 1 Infection. J. Virol. 92 (2018).

24. K. M. Whitworth, et al., Gene-edited pigs are protected from porcine reproductive and respiratory syndrome virus. Nat. Biotechnol. 34, 20–22 (2016).

25. J. Chen, et al., Generation of pigs resistant to highly pathogenic-porcine reproductive and respiratory syndrome virus through gene editing of CD163. Int. J. Biol. Sci. 15, 481–492 (2019).

26. R. M. Anderson, The concept of herd immunity and the design of community-based immunization programmes. Vaccine 10, 928–935 (1992).

27. R. M. Anderson, R. M. May, Infectious Diseases of Humans: Dynamics and Control (Oxford University Press, 1992).

28. E. Pileri, et al., Vaccination with a genotype 1 modified live vaccine against porcine reproductive and respiratory syndrome virus significantly reduces viremia, viral shedding and transmission of the virus in a quasi-natural experimental model. Vet. Microbiol. 175, 7–16 (2015).

29. G. Nodelijk, et al., Introduction, persistence and fade-out of porcine reproductive and respiratory syndrome virus in a Dutch breeding herd: a mathematical analysis. Epidemiol. Infect. 124, 173– 182 (2000).

30. C. Burkard, et al., Precision engineering for PRRSV resistance in pigs: Macrophages from genome edited pigs lacking CD163 SRCR5 domain are fully resistant to both PRRSV genotypes while maintaining biological function. PLoS Pathog. 13, 1–28 (2017).

31. V. Bitsouni, S. Lycett, T. Opriessnig, A. Doeschl-Wilson, Predicting vaccine effectiveness in livestock populations: A theoretical framework applied to PRRS virus infections in pigs. PLoS One 14, e0220738 (2019).

32. O. C. Ezezika, et al., Factors influencing agbiotech adoption and development in sub-Saharan Africa. Nat. Biotechnol. 30, 38–40 (2012).

33. M. Amadori, E. Razzuoli, Immune Control of PRRS: Lessons to be Learned and Possible Ways Forward. Front. Vet. Sci. 1, 1–14 (2014).

34. M. J. Keeling, B. T. Grenfell, Individual-based perspectives on R0. J. Theor. Biol. 203, 51–61 (2000).

35. T. Britton, Epidemics in heterogeneous communities: Estimation of R0 and secure vaccination coverage. J. R. Stat. Soc. Ser. B Stat. Methodol. 63, 705–715 (2001).

36. C. A. Corzo, et al., Control and elimination of porcine reproductive and respiratory syndrome virus. Virus Res. 154, 185–192 (2010).

37. R. R. R. Rowland, J. Lunney, J. Dekkers, Control of porcine reproductive and respiratory syndrome (PRRS) through genetic improvements in disease resistance and tolerance. Front. Genet. 3, 1–6 (2012).

38. A. R. Allen, R. A. Skuce, A. W. Byrne, Bovine tuberculosis in Britain and Ireland - A perfect storm? The confluence of potential ecological and epidemiological impediments to controlling a chronic infectious disease. Front. Vet. Sci. 5, 1–17 (2018).

39. X. J. Meng, Heterogeneity of porcine reproductive and respiratory syndrome virus: Implications for current vaccine efficacy and future vaccine development. Vet. Microbiol. 74, 309–329 (2000).

40. E. Mateu, I. Diaz, The challenge of PRRS immunology. Vet. J. 177, 345–351 (2008).

41. G. Reiner, Genetic resistance - an alternative for controlling PRRS? Porc. Heal. Manag. 2, 27 (2016).

42. J. R. Dunkelberger, P. K. Mathur, M. S. Lopes, E. F. Knol, J. C. M. Dekkers, “Pigs Can Be Selected forIncreased Natural Resistance to PRRS Without Affecting Overall Economic Value in the Absence ofPRRS” (2017).

43. E. Pileri, E. Mateu, Review on the transmission porcine reproductive and respiratory syndrome virus between pigs and farms and impact on vaccination. Vet. Res. 47, 1–13 (2016).

44. A. S. Fahrion, E. grosse Beilage, H. Nathues, S. Dürr, M. G. Doherr, Evaluating perspectives for PRRS virus elimination from pig dense areas with a risk factor based herd index. Prev. Vet. Med. 114, 247–258 (2014).

45. K. Hanada, Y. Suzuki, T. Nakane, O. Hirose, T. Gojobori, The origin and evolution of porcine reproductive and respiratory syndrome viruses. Mol. Biol. Evol. 22, 1024–1031 (2005).

46. Y. Nan, et al., Improved vaccine against PRRSV: Current Progress and future perspective. Front. Microbiol. 8, 1–17 (2017).

47. P. H. Rathkjen, J. Dall, Control and eradication of porcine reproductive and respiratory syndrome virus type 2 using a modified-live type 2 vaccine in combination with a load, close, homogenise model: an area elimination study. Acta Vet. Scand. 59, 4 (2017).

48. P. Visscher, R. Pong-Wong, C. Whittemore, C. Haley, Impact of biotechnology on (cross)breeding programmes in pigs. Livest. Prod. Sci. 65, 57–70 (2000).

49. J. C. M. Dekkers, P. M. Mathur, E. J. Huff-Lonergan, “Genetic improvement of the pig” in The Genetics of the Pig, 2nd Edition, M. Rothschild, A. Ruvinsky, Eds. (CABI, 2011), pp. 390–425.

50. P. Gottardo, et al., A strategy to exploit surrogate sire technology in livestock breeding programs. G3 Genes, Genomes, Genet. 9, 203–215 (2019).

51. S. Gonen, et al., Potential of gene drives with genome editing to increase genetic gain in livestock breeding programs. Genet. Sel. Evol. 49, 1–14 (2017).

52. C. Noble, et al., Daisy-chain gene drives for the alteration of local populations. Proc. Natl. Acad. Sci. U. S. A. 116, 8275–8282 (2019).

53. A. L. Norris, et al., Template plasmid integration in germline genome-edited cattle in BioRxiv, (2019), p. 715482.

54. Pig333 Professional Pig Community, Pig Production Data - Annual Pig Census.

55. W. Hall, E. Neumann, Fresh Pork and Porcine Reproductive and Respiratory Syndrome Virus: Factors Related to the Risk of Disease Transmission. Transbound. Emerg. Dis. 62, 350–366 (2015).

56. H. McCallum, N. Barlow, J. Hone, How should pathogen transmission be modelled? Trends Ecol. Evol. 16, 295–300 (2001).

57. A. B. Doeschl-Wilson, et al., Combining laboratory and mathematical models to infer mechanisms underlying kinetic changes in macrophage susceptibility to an RNA virus. BMC Syst. Biol. 10, 101 (2016).

58. K. Oldenbroek, L. van der Waaij, Textbook Animal Breeding and Genetics for BSc Students (Centre for Genetic Resources The Netherlands and Animal Breeding and Genomic Centre, 2015).

59. T. Hai, F. Teng, R. Guo, W. Li, Q. Zhou, One-step generation of knockout pigs by zygote injection of CRISPR/Cas system. Cell Res. 24, 372–375 (2014).

