## Supplemental Figures and Tables for "Gene editing in Farm Animals: A Step Change for Eliminating Epidemics on our Doorstep?"

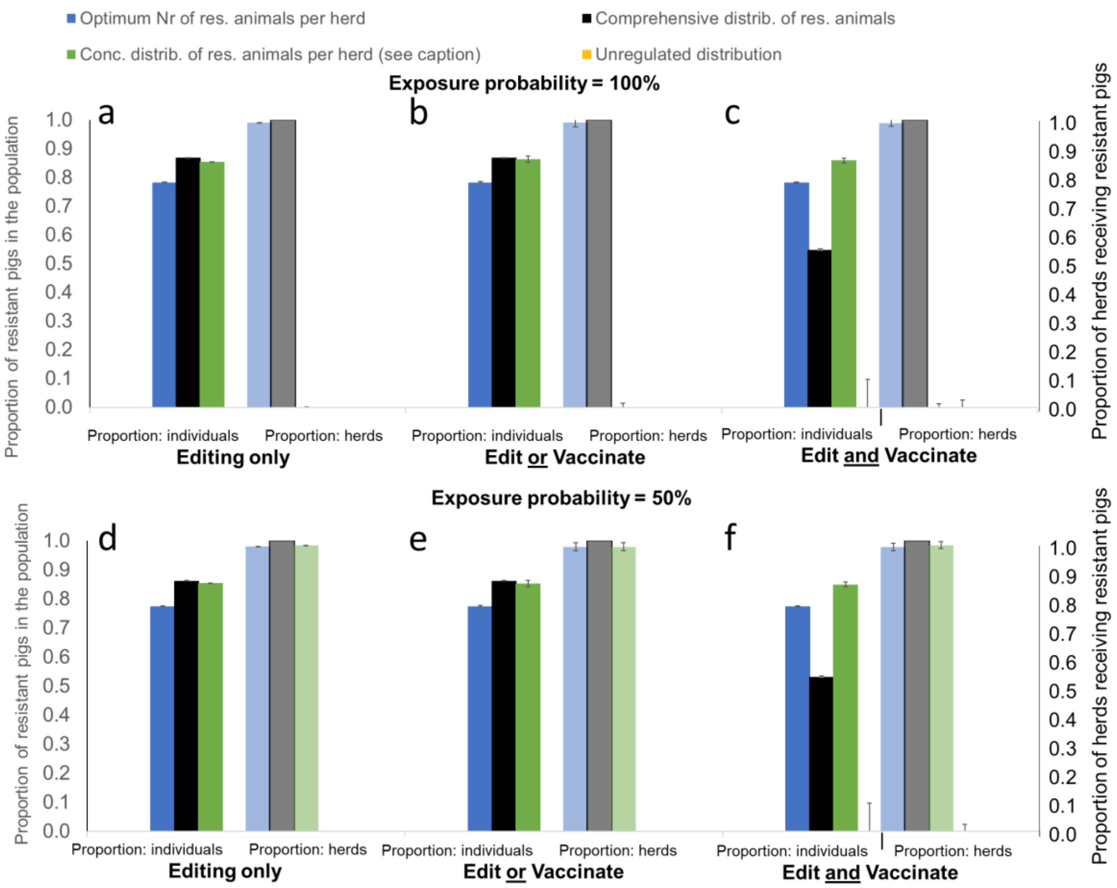


**Fig. S1.** Minimum required proportion of genetically resistant animals (solid bars) and corresponding herds adopting gene editing (transparent bars) for achieving disease elimination through gene editing alone or with vaccination combined, depending on how edited animals are distributed across the herds. Results are shown for average $R_0$ value of 5 and exposure probability of either 100% (Fig.2 a-c) and 50% (Fig.2 d-f), and vaccine effectiveness of 70%. Different colours refer to different distribution scenarios (see Table 1) with blue = Optimum, black = Comprehensive, green = Concentrated and yellow = Unregulated (not depicted here as elimination was not feasible)

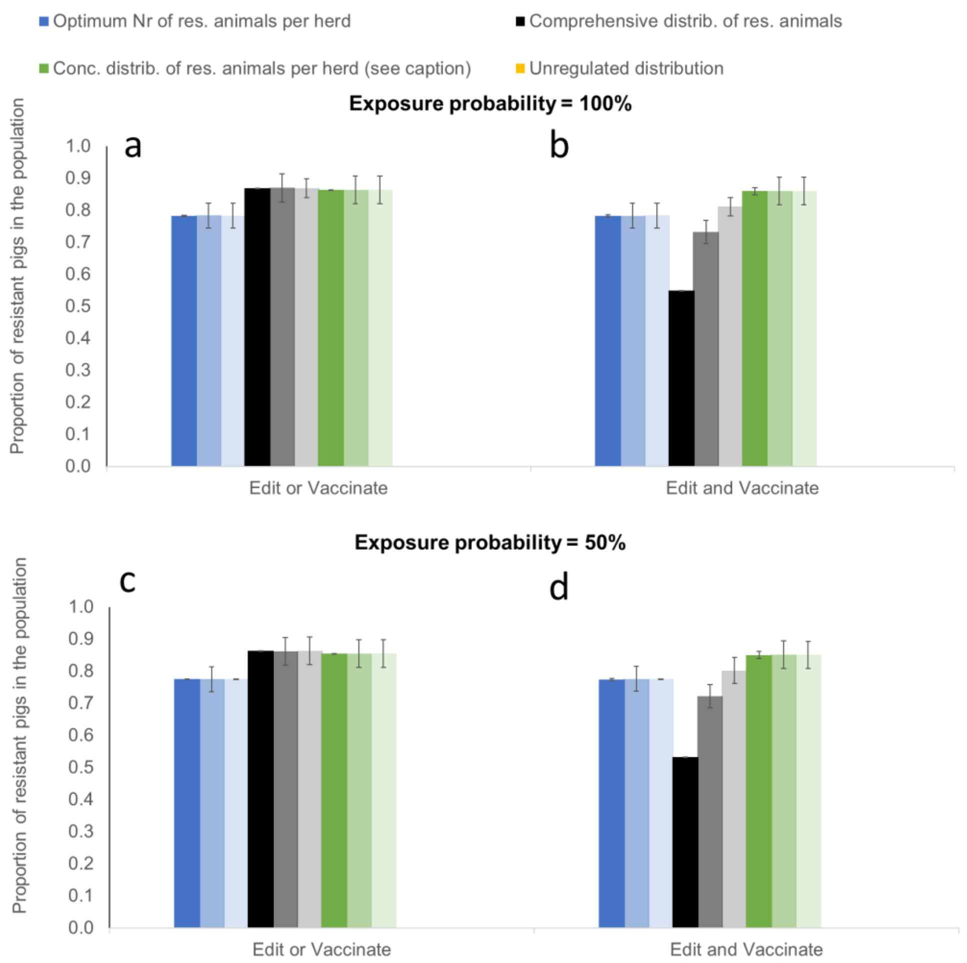


**Fig. S2**. Minimum required proportion of genetically resistant animals for achieving disease elimination through gene editing and vaccination combined, depending on vaccine effectiveness $\varepsilon_V$ and exposure probability. Solid bars: $\varepsilon_V$ = 0.7, 50% transparency bars: $\varepsilon_V$ = 0.5; 80% transparency bars: $\varepsilon_V$ = 0.3. Different colours refer to different distribution scenarios with blue = Optimum, black = Comprehensive, green = Concentrated and yellow = Unregulated (not depicted here as elimination was not feasible). An average transmission potential of $R_0$ = 5 was assumed.

**Table S1.** Time in months to reach the proportion of genetically resistant pigs in the commercial population required for PRRS elimination under different elimination strategies: Presented results correspond to the distribution strategies associated with the minimum / maximum proportion of genetically resistant pigs required to achieve elimination in the case of *Gene Editing Only* without use of vaccination, with complementary vaccination in herds not receiving resistant pigs only (*Edit or Vaccinate*), and complementary vaccination of all susceptible animals (*Edit and Vaccinate*), respectively. An average $R_0$ of 1.5 and 100% exposure probability was assumed.

| | | **Proportion of pigs selected for editing** | | |
|---|---|---|---|---|
| | | **20%** | **10%** | **5%** |
| | Number of edits before 100% of pigs are resistant: | 24798 | 12637 | 6571 |
| **Scenario** | | **Time (months)** | | |
| **Editing only – compreh.** | **74%** resistant commercial pigs reached after: | 61 | 65 | 67 |
| **– optimal** | **30%** resistant commercial pigs reached after: | 43 | 47 | 51 |
| **Edit or Vac – compreh.** | **74%** resistant commercial pigs reached after: | 61 | 65 | 67 |
| **– optimal** | **21%** resistant commercial pigs reached after: | 39 | 44 | 47 |
| **Edit and Vac – compreh.** | **12%** resistant commercial pigs reached after: | 34 | 38 | 42 |
| **– unregulated** | **70 %** resistant commercial pigs reached after: | 59 | 63 | 65 |

**Table S2.** List of input parameters and their assumed values for the epidemiological model.

| **Parameter** | **Description** | **Assumed value(s)** |
|---|---|---|
| N | Total number of pigs in national population | 12 Million (1) |
| $n_H$ | Number of herds | 5,000 (1) |
| $\mu_H$, $\sigma_H$ | Average herd size and standard deviation | $\mu_H$ = 2,400 ; $\sigma_H$ = 1,000 (2) |
| $\mu_{R0}$, $\sigma_{R0}$ | Mean value and standard deviation, respectively for the basic reproductive ratio $R_0$ across all herds | $\mu_{R0}$ was varied between 1.1 and 5 (3, 4); $\sigma_{R0}$ = 1 |
| $\varepsilon_e$ | Efficacy of gene editing | 1 (5–7) |
| $\varepsilon_v$ | Vaccine effectiveness | Varied between 0.3 and 0.7§1 (8–10) |
| $P_e$ | Proportion of genetically resistant pigs in a herd | either assumed equal in all herds with a fixed value of 0.1, 0.5 or $\left(1 - 1/(\mu_{R_0} + 2.56\sigma_{R_0})\right)$ §2 or set to the herd specific critical value $P_e^*$ defined in equation [2]. |
| $P_v$ | Proportion of vaccinated pigs in a herd | Varied between 0 and 1, depending on the simulated scenario |
| $p_{exp}$ | Exposure probability | 0.5 or 1 |

§1 Vaccine effectiveness $\varepsilon_v \leq 0.7$ were chosen as no PRRS vaccine to date is fully protective against infection with all circulating PRRSv strains. The values imply that PRRS cannot be eliminated by vaccination alone.

§2 This value corresponds to the minimum fixed proportion of edits required per herd for achieving $R < 1$ in 99% of herds, as per eq. [2]. It refers to the more realistic situation where the distribution parameters $\mu_{R0}$, $\sigma_{R0}$ rather than the herd-specific R0-values are assumed known. Note that for $\mu_{R0}$ =1.5, this value is ~0.75.

**Table S3.** Initial selection proportions of individuals selected in the different tiers of the breeding pyramid.[1]

| **Classes** | **Tier** | **Selection proportion** |
|---|---|---|
| SPF nucleus males mated to SPF nucleus females | I | 0.02 |
| First parity nucleus gilts used within SPF | I | 0.10 |
| SPF gilts transferred to production nucleus | II | 0.40 |
| SPF semen transferred to production nucleus | II | 0.10 |
| Production nucleus gilts retained for use | II | 0.20 |
| SPF nucleus semen transferred to multiplier | III | 0.10 |
| Production nucleus gilts transferred to multiplier | III | 0.50 |
| F1 gilts from tier III transferred to breeder weaner herds | IV | 0.60 |
| SPF semen transferred to breeder weaner herds | IV | 0.10 |

**Table S4.** Assumed values for reproduction and live cycle parameters applied in the pig breeding pyramid simulation model. Source (11)

| **Parameter** | **Value** |
|---|---|
| Sow gestation length | 4 months |
| Farrowing interval | 5 months |
| Gilt age at first mating | 8 months |
| Boar age at first mating / semen provision | 8 months |
| Litter size (No of piglets) | 12 |
| Maximum parities per sow (= culling age in years) | 8 |
| Maximum age of boars at provision of semen (= culling age in years) | 4 |

**References**


1. Pig333 Professional Pig Community, Pig Production Data - Annual Pig Census.
2. Food and Agriculture Organization of the United Nations, FAOSTAT Database (2019) (March 14, 2019).
3. G. Nodelijk, *et al.*, Introduction, persistence and fade-out of porcine reproductive and respiratory syndrome virus in a Dutch breeding herd: a mathematical analysis. *Epidemiol. Infect.* **124**, 173–182 (2000).
4. C. Charpin, *et al.*, Infectiousness of pigs infected by the Porcine Reproductive and Respiratory Syndrome virus (PRRSV) is time-dependent. *Vet. Res.* **43** (2012).
5. K. M. Whitworth, *et al.*, Gene-edited pigs are protected from porcine reproductive and respiratory syndrome virus. *Nat. Biotechnol.* **34**, 20–22 (2016).
6. C. Burkard, *et al.*, Precision engineering for PRRSV resistance in pigs: Macrophages from genome edited pigs lacking CD163 SRCR5 domain are fully resistant to both PRRSV genotypes while maintaining biological function. *PLoS Pathog.* **13**, 1–28 (2017).
7. C. Burkard, *et al.*, Pigs Lacking the Scavenger Receptor Cysteine-Rich Domain 5 of CD163 Are Resistant to Porcine Reproductive and Respiratory Syndrome Virus 1 Infection. *J. Virol.* **92** (2018).
8. W. L. Mengeling, K. M. Lager, A. C. Vorwald, D. F. Clouser, Comparative safety and efficacy of attenuated single-strain and multi-strain vaccines for porcine reproductive and respiratory syndrome. *Vet. Microbiol.* **93**, 25–38 (2003).


[1] These parameters are used as reference and represent common practices in most breeding schemes [personal communication Dr. Peter Amer]

9. F. A. Zuckermann, *et al.*, Assessment of the efficacy of commercial porcine reproductive and respiratory syndrome virus (PRRSV) vaccines based on measurement of serologic response, frequency of gamma-IFN-producing cells and virological parameters of protection upon challenge. *Vet. Microbiol.* **123**, 69–85 (2007).
10. V. G. Papatsiros, Porcine respiratory and reproductive syndrome virus vaccinology: A review for commercial vaccines. *Am. J. Anim. Vet. Sci.* **7**, 149–158 (2012).
11. I. Kyriazakis, C. T. Whittemore, *Whittemore's Science and Practice of Pig Production*, Third Edit (Blackwell Publishing, 2006).
